# Detecting cell-level transcriptomic changes of Perturb-seq using Contrastive Fine-tuning of Single-Cell Foundation Models

**DOI:** 10.1101/2025.04.17.649395

**Authors:** Wenmin Zhao, Ana Solaguren-Beascoa, Grant Neilson, Regina Reynolds, Louwai Muhammed, Liisi Laaniste, Sera Aylin Cakiroglu

**Affiliations:** Cosyne Therapeutics, London, UK

## Abstract

Genome-scale perturbation cell atlases are an exciting new resource to understand the transcriptomic and phenotypic impact of single-gene activation or knockdown. However, in terms of differentially expressed genes identified, the signal detected in these data atlases is low, leading to the exclusion of most data from downstream analyses. Recent advances in single-cell foundation models have shown promise in capturing complex biological insights. However, their application to perturbation analysis, especially in predicting perturbed single-cell transcriptomes, remains limited. In this paper, we focus on learning representations of single-cell transcriptomes that capture subtle, yet important, transcriptome-wide changes, and we propose a novel fine-tuning strategy using contrastive learning to leverage single-cell foundation models for this task. We pre-train a single-cell foundation model and fine-tune on a genome-scale perturbation dataset using a contrastive loss, which minimises the distance between cell embeddings from unperturbed cells while maximising the distance between perturbed and unperturbed cells. We validate and test the model on unseen perturbations, demonstrating its ability to identify global biologically meaningful transcriptional changes not captured by traditional differential expression methods. Our approach provides a novel framework for analysing single-cell perturbation data and offers a more effective means of identifying perturbations that drive systemic gene expression changes.

## 1 Introduction

Understanding the transcriptomic and phenotypic outcomes of perturbed gene expression (e.g. knockdown or activation) at the single-cell level has the potential to improve our understanding of development, disease mechanisms, and cell-state engineering, with applications in target identification for drug discovery. High-throughput methods based on CRISPRi perturbations have recently been applied to this end at genome scale: Replogle et al. (2022) perturbed just under 10k expressed genes, resulting in a dataset of close to 2M single-cell transcriptomes and new preprints signpost genome-scale perturbation cell atlases as a new frontier of single-cell data (Nourreddine et al., 2024). Simultaneously, advanced deep-learning techniques have been applied in the prediction of perturbed single-cell transcriptome data (Roohani et al., 2023; Hao et al., 2024). Recent developments in training so-called ‘single-cell foundation models’ on large-scale single-cell transcriptomic data (Theodoris et al., 2023; Yang et al., 2022; Cui et al., 2023; Hao et al., 2024), have renewed excitement about learning fundamental representations of biology and their application to understanding co-regulatory effects. However, on the task of predicting gene expression under perturbation, these models have been outperformed by simple baselines (Bendidi et al., 2024; Ahlmann-Eltze et al., 2024a; Gaudelet et al., 2024).

The main objective of perturbation studies is to identify genes that drive meaningful transcriptional changes. Usually, a transcriptomic perturbation signature is defined by identifying genes exhibiting the most significant changes in expression upon perturbation of the target gene. However, differential expression analyses often treat each gene in isolation, failing to account for the intricate interdependencies between genes that are crucial to understanding perturbation effects at the cellular level. This approach also has unique challenges when applied to single-cell perturbation data, due to high sparsity (e.g. zero-inflated gene expression distributions), as well as varying perturbation efficiency of single-guide RNAs (sgRNAs), which is difficult to distinguish from biological signal or technical noise. Together with a limited number of cells per perturbation, this results in difficulties in identifying differentially expressed genes in 70-89% of the genome-wide perturbations performed (Replogle et al., 2022; Nourreddine et al., 2024). It is unclear how many of these perturbations give rise to a meaningful transcriptomic signal, and they are commonly excluded from any downstream analyses. To fully utilise genome-scale perturbation data atlases, approaches need to identify perturbations that elicit a change in the overall transcriptomic state of the cell instead of focusing only on differentially expressed genes.

In this paper, we introduce a method for learning representations of single-cell transcriptomes that encode information about the perturbation state (perturbed/unperturbed) based on a cell’s whole transcriptome. To this end, we present a novel fine-tuning strategy for single-cell foundation models using contrastive learning. By fine-tuning these models on single-cell perturbation data, we aim to improve their ability to capture and predict perturbation-driven transcriptomic changes even when no differentially expressed genes can be detected. We show that our approach captures more signals in the data than previous methods, allowing for a more comprehensive use of perturbation analysis in single-cell biology.

## 2 Related Work

### 2.1 Perturbation Analysis

There is no standard analysis for identifying perturbations that induce transcriptomic changes in single-cell RNA sequencing (scRNA-seq) data. Most approaches rely on differential gene expression analysis (Replogle et al., 2022; Nourreddine et al., 2024), using statistical tests like the Mann-Whitney U test or regression models fitted to expression values with a negative binomial distribution (Alessandrì et al., 2019; Chen et al., 2025). A drawback of these methods is the focus on large individual gene-level changes to determine the impact of perturbations, which may miss larger global shifts created by small combinatorial changes. Approaches that aim to determine distribution shifts throughout the transcriptome use dimensionality reduction with Principal Component Analysis (PCA), followed by computing distance metrics such as the energy distance (Peidli et al., 2024). Other approaches that are commonly used to project high-dimensional scRNA-seq data into lower-dimensional embeddings include Variational Autoencoders (VAEs), exemplified by models such as scVI (Lopez et al., 2018), which have recently been compared in benchmarks of perturbation analyses (Bendidi et al., 2024). Recently published single-cell foundation models, such as Geneformer (Theodoris et al., 2023) and scGPT (Cui et al., 2023), are transformer-based models that are pretrained on large-scale single-cell atlases and then fine-tuned on a range of downstream tasks, where they often outperform existing methods. For perturbation analyses, however, these models are still outperformed by simple methods like PCA (Bendidi et al., 2024; Ahlmann-Eltze et al., 2024b).

### 2.2 Contrastive Learning

Contrastive learning is an approach that helps to extract meaningful representations by contrasting similar and dissimilar pairs of data points; it leverages the assumption that similar examples should be closer in a learned embedding space, while dissimilar examples should be farther apart. This type of training is commonly used in image-caption pre-training and fine-tuning (Radford et al., 2021; Jia et al., 2021; Zhai et al., 2022; 2023), where aligning image and text representations in a shared embedding space enables good performance on zero-shot transfer tasks, such as classification and retrieval. Recently, there has also been a surge in the use of contrastive learning to fine-tune encoder-only language models for improved sentence representation learning (Gao et al., 2021; Zhang et al., 2022; Chuang et al., 2022). In particular, contrastive learning can alleviate the problem of anisotropy, where sentence embeddings occupy only a narrow cone in the embedding space, leading to poor sentence representation and low performance in downstream tasks (Xu et al., 2023).

## 3 Methods

### 3.1 Pre-training ON Large Scale Single-Cell RNA-SEQ Data

We downloaded scRNA-seq data from ∼ 33*M* unique cells across 265 datasets in the census dataset (version 2023-07-25) from the CellXGene data portal (CZI Single-Cell Biology Program et al., 2023). Data processing followed Theodoris et al. (2023), representing each single-cell transcriptome as a sequence of gene names of maximum length 2, 048 ordered by their median-normalised expression. We excluded cancer cells and cells with *<* 500 expressed genes. Our single-cell foundation model is based on a bidirectional transformer encoder-only architecture (BERT) similar to Geneformer (Theodoris et al., 2023), receiving a single-cell transcriptome as an ordered sequence of gene names of maximum length 2, 048. The model was pre-trained with a masked language modelling task (masking 15% of input tokens) for three epochs. For more implementation details see Appendix A.1, and a schematic of the model is shown in Figure 1A. We compared our model and Geneformer on a dataset that was recently published and, therefore, did not form part of the training data for either model (Heimlich et al., 2024). The two models performed comparably, with our model slightly outperforming Geneformer in reproducing the overall ranking of highly expressed genes (Appendix A.2 and Supplementary Figure A.1).

**Figure 1.**
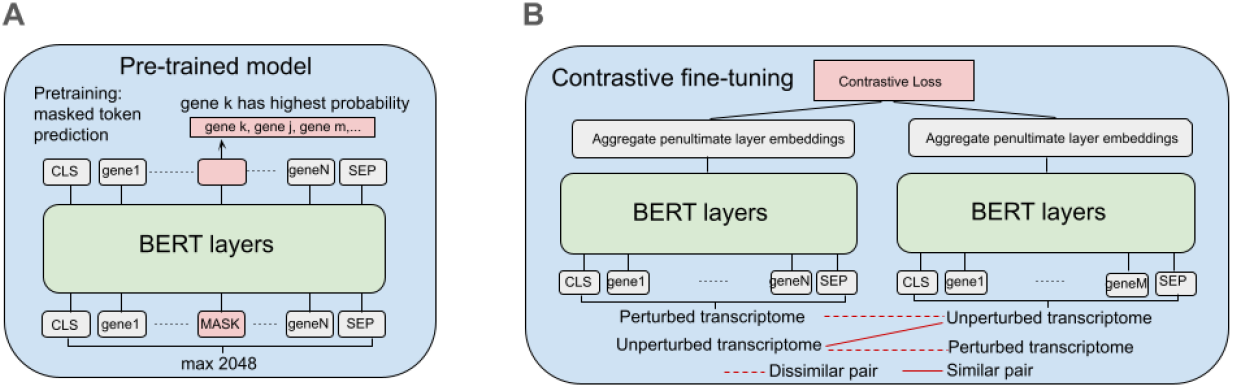
A: Schematic of the pre-trained foundation model. The model is trained with a masked language modelling task where input tokens are randomly masked and the model is trained to predict the right gene name. B: Schematic of the model fine-tuning task. The input to the model is a pair of transcriptomes: a perturbed and unperturbed cell form a dissimilar pair, or two unperturbed cells are a similar pair. The gene embeddings of the penultimate layer of the pre-trained model are aggregated into a cell embedding and the contrastive loss shown in Equation 1 is calculated for embedding pairs in each batch and back-propagated through all layers of the model.

### 3.1 Contrastive Fine-tuning on Perturb-seq

We leveraged the largest genome-scale perturbation dataset to date of 9,866 perturbations (knockdowns) in 1.98 million K562 lymphoblast cells (Replogle et al., 2022). For each cell, we recorded the perturbed gene or noted it as “unperturbed” if a non-targeting sgRNA was used. There are 75, 000 unperturbed cells, and the median number of cells per perturbation is ∼ 200 (Figure A.2). For each perturbation, we identified differentially expressed genes (DEGs) between perturbed and unperturbed cells using a Mann-Whitney U test (Supplementary Figure A.3). Using the E-distance between perturbed and unperturbed PCA embeddings as described in Section 3.4, we observed a Pearson correlation of 0.54 between E-distance and DEG counts. We selected 1, 541 perturbations with ≥ 20 DEGs for fine-tuning, as these are more likely to exhibit larger overall transcriptional changes, resulting in a stronger training signal. In our evaluation, however, we assessed the model’s generalisation to putative perturbations for which no DEGs were detected; that is, perturbations that induce subtle yet widespread transcriptional changes, even when the expression of individual genes do not show large shifts. Perturb-seq data was normalised with gene medians from the pre-training dataset before ranking genes by their expression into an ordered sequence of gene names, as described above. We grouped perturbed cells based on their perturbed target gene, and split cells by target gene into train, validation and test sets by a 80/10/10 rule. We split the unperturbed cells using the same 80/10/10 rule, and randomly sampled size-matched sets from the training set for each training perturbation to provide examples of control/dissimilar cells during training.

We used the final checkpoint of our pre-trained model for fine-tuning. A single training sample consisted of a pair of single-cell transcriptomes that are either “similar” or “dissimilar”: a perturbed and unperturbed cell constituted a dissimilar pair, whereas two unperturbed cells formed a similar pair. The cell embeddings for both transcriptomes in the pair were obtained from the model as described in Section 3.3. The following contrastive loss was calculated for embedding pairs in each batch and back-propagated through all layers of the model:

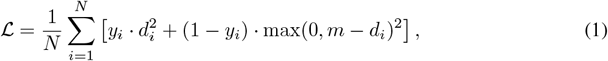

where *N* is the batch size, *m* is the margin hyperparameter, and 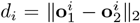 is the Euclidean distance between the two embedding vectors **o**^*i*^ and **o**^*i*^ of the single-cell transcriptomes belonging to the *i*th pair in the batch, and *y*_*i*_ ∈ {0, 1} is the corresponding binary label, where 1 indicates that the pair is similar, and 0 dissimilar. The term max(0, *m* − *d*_*i*_) ensures that the dissimilar loss is non-negative and only contributes when *d*_*i*_ is less than the margin. A schematic of the contrastive fine-tuned model is shown in Figure 1.

We ensured that cells with the same perturbed gene were in the same batch when training on dissimilar pairs, alternating between training on batches of similar and dissimilar pairs. Note that we did not explicitly train the model to learn separations between perturbations. We fine-tuned the models using a margin of 20, a learning rate of 1*e* − 5, using the AdamW optimiser with a weight decay of 1*e* − 2. Fine-tuning of 10 epochs took 7 days on a single NVIDIA T4 15G GPU.

### 3.3 Cell embeddings

To obtain cell embeddings from the pre-trained or fine-tuned model, we performed a forward pass on an input transcriptome and extracted the contextualised embedding for each input gene from the penultimate layer. To compute the cell embedding, we took the mean of all of the cell’s gene embeddings to form a single embedding vector of dimension 256, similar to Theodoris et al. (2023).

### 3.4 Distance Metrics

Our objective is to learn embeddings of single-cell transcriptomes that capture transcriptomic differences between perturbed and unperturbed cells. To evaluate our model, we used a set of metrics designed to measure how well the embeddings separated perturbed from unperturbed cells, as well as different pairs of perturbations. Although the model was not directly trained to do the latter, this evaluation tests whether the model captures more general differences in transcriptomic states. To measure the quality of the embeddings in this regard, we used the following metrics:

- **Energy distance (E-distance)**: This metric measures the statistical dispersion between two groups, making it particularly useful for quantifying the separation between distributions. Here, it captures the difference in expression profiles between perturbed and unperturbed cells, taking into account the variation within the two groups (Peidli et al., 2024). Higher values indicate greater separation, suggesting that a model is better at distinguishing transcriptional changes caused by perturbations. The distance between the distributions *P* and *Q* is defined as

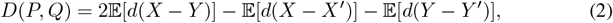

where, *X, X*^*′*^ ∼ *P* and *Y, Y* ^*′*^ ∼ *Q* are samples drawn from *P* and *Q*, respectively, and *d* is the Euclidean distance. When calculating *D* for pairs of perturbations, *X* and *Y* are the whole sets of the cell embeddings for each perturbation target, whereas when computing *D* for perturbed and unperturbed cells, *Y* is replaced by embeddings of random samples from the unperturbed cell population (size-matching *X*).
- **Cosine E-distance:** Similar to E-distance, but *d* in Equation 2 is taken to be the cosine distance instead of the Euclidean distance. This change makes the distance invariant to the magnitudes of the embedding vectors that are compared.
- **High-dimensional Wasserstein distance:** This metric quantifies the minimal effort required to transform one distribution into another. While the E-distance emphasizes group separation, this metric provides a more nuanced measure of difference in distributions.

To compare the separation quality of different model embeddings, we L2-normalised each embedding to account for variations in embedding dimensions. We then applied a z-score normalisation to each metric using a control distribution, which was derived from pairs of embedding groups randomly sampled from the unperturbed cell population, to account for different variability and scales in the embeddings of the unperturbed distribution. The normalisation of metric *D* is given by

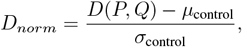

where *µ*_control_ and *σ*_control_ denote the mean and standard deviation of the control distribution, and *D*_*norm*_ is the normalised distance.

## 4 Results

### 4.1 Contrastive fine-tuning allows the encoding of distinct transcriptomic states in cell embeddings

For a visual assessment of the impact of contrastive fine-tuning, Figure 2 shows UMAP projections of embeddings of perturbed and unperturbed cells in the test set from the pre-trained and the fine-tuned model. Before fine-tuning, there is significant mixing between the embeddings of the perturbed and unperturbed cells, indicating that the pre-trained model is unable to distinguish between them. There is also a separate cluster that contains a mixture of perturbed and unperturbed cells, likely capturing batch effects. After contrastive fine-tuning, however, the model’s embeddings separate better between perturbed and unperturbed cells: embeddings of the unperturbed cells now cluster together, and no strong batch effects are apparent. While there is some overlap of unperturbed and perturbed cell embeddings, there is a more pronounced distinction between them, suggesting that the fine-tuning process has improved the model’s ability to detect transcriptomic changes.

**Figure 2.**
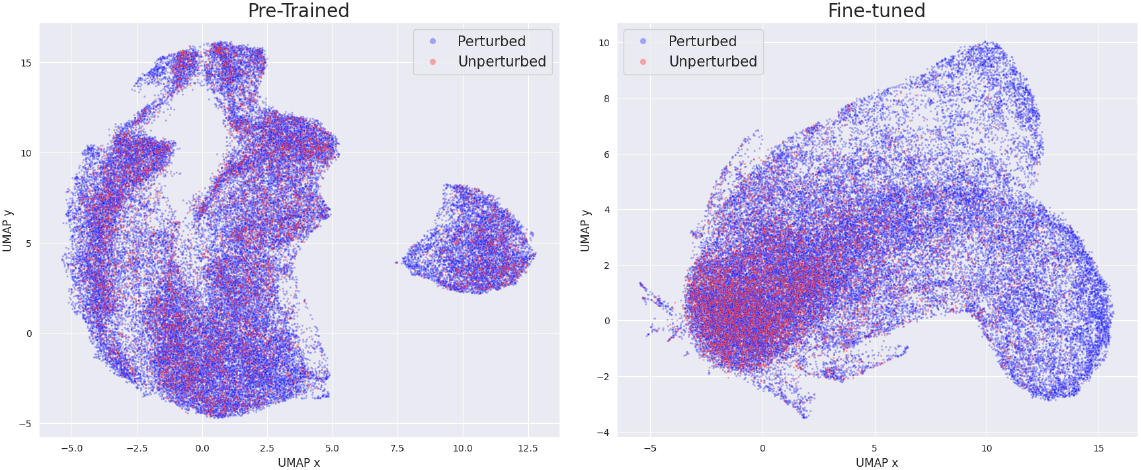
UMAP visualisation of cell embeddings in the test set from the pre-trained model before (left) and after contrastive fine-tuning (right). Perturbed cells (*N* = 47, 349) of 232 perturbations are shown in blue, and unperturbed cells (*N* = 7, 470) in red.

Figure 3 shows examples of cell embeddings for two perturbations, *TAF10* and *DHDDS*, and their separation from a size-matched random sample of unperturbed cells from the test set. These perturbations rank among the top five in the test set with the largest E-distance in the pre-trained model. In both cases, cell embeddings of perturbed cells separate from embeddings of unperturbed cells after contrastive fine-tuning, indicating the model’s increased ability to distinguish cells with different transcriptomic states.

**Figure 3.**
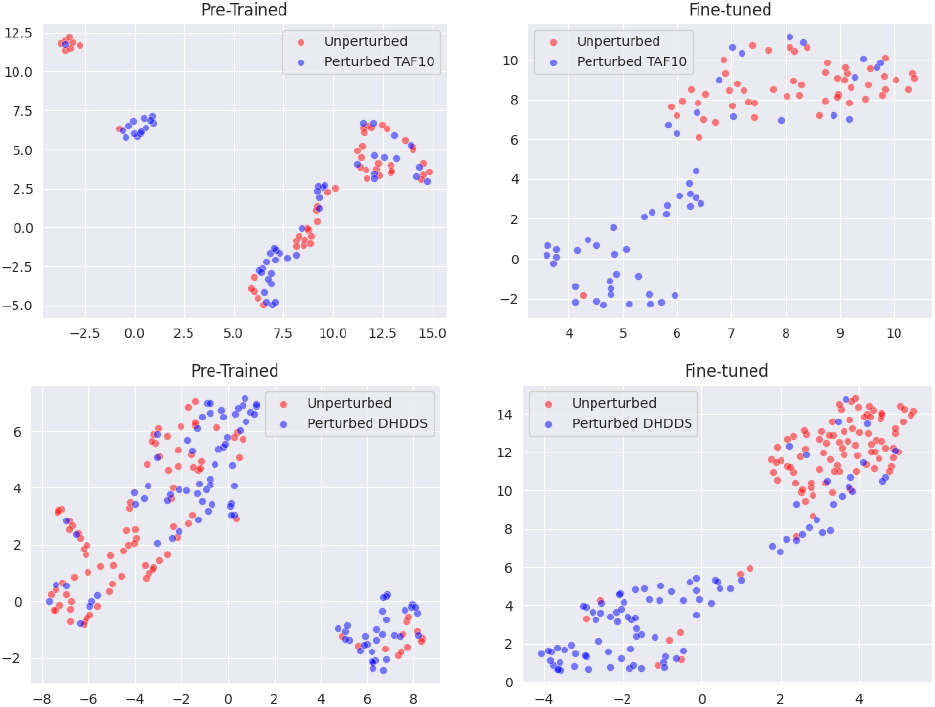
UMAP visualisation of embeddings of perturbed (blue) and unperturbed (red) cells from the test set for the perturbation targets *TAF10* (top) and *DHDDS* (bottom), before (left) and after contrastive fine-tuning (right).

While the model was not explicitly trained to distinguish transcriptomes from different perturbation targets, we hypothesised that if the model uses transcriptomic states to distinguish unperturbed and perturbed cells, then it should capture these changes also for different perturbation targets. Figure 4 shows UMAP visualisations of cell embeddings before and after contrastive fine-tuning for the perturbation targets *LARP7* vs *MRPS33* and *PCBP2* vs *POLRMT*. In both cases, the fine-tuned model is able to distinguish between transcriptomes arising from the different unseen perturbations, further showcasing the model’s ability to capture more nuanced perturbation effects after contrastive fine-tuning.

**Figure 4.**
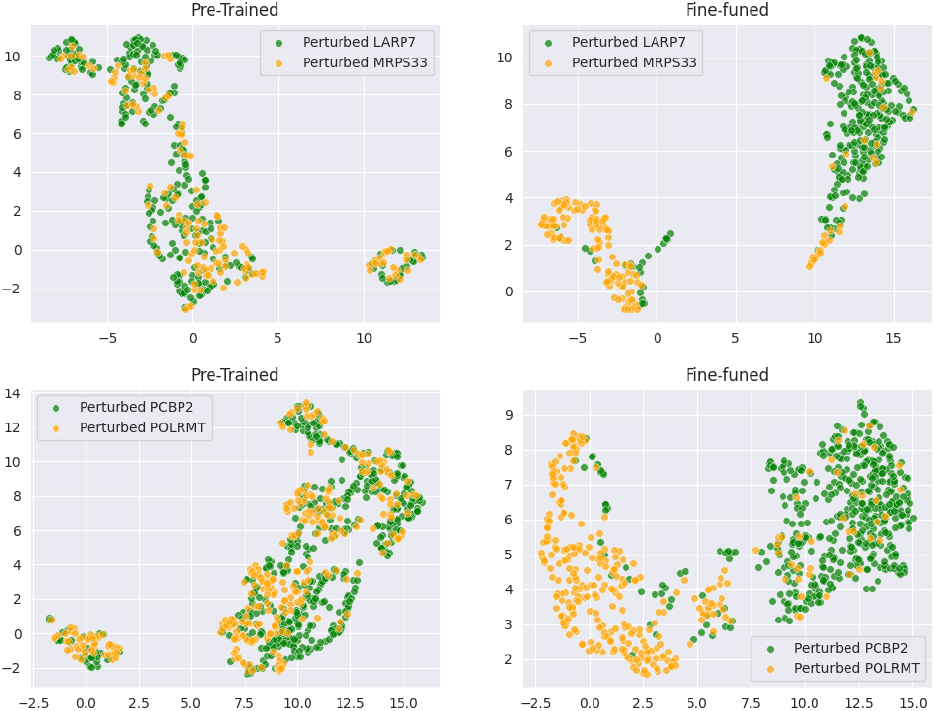
UMAP visualisation of embeddings of perturbed cells from the test set with perturbation targets *LARP7* vs *MRPS33* (top) and *PCBP2* vs *POLRMT* (bottom), before (left) and after contrastive fine-tuning (right).

### 4.2 Contrastive fine-tuning outperforms strong baseline models

To assess the ability of our fine-tuned model to capture transcriptomic states upon perturbation, we quantified the model’s performance with three distance metrics described in Section 3.4. We compared our model against five other approaches, some of which have been shown to outperform deep learning models in perturbation analysis (Bendidi et al., 2024):

1. Pre-trained: the single-cell foundation model before fine-tuning.
2. Highly variable genes (HV): using the expression of the top 300 highly variable genes across the whole dataset (including both perturbed and unperturbed cells).
3. PCA: applying PCA on the whole dataset (log normalised) and selecting the top 200 principal components.
4. scVI Linear: a variational autoencoder with a single encoder and decoder layer trained on raw expression counts of all cells in the training set of the contrastive model (Lopez et al., 2018). See A.3 for training and implementation details.
5. scVI 5L: scVI where both the encoder and decoder contain 5 layers trained on raw expression counts of all cells in the training set of the contrastive model (Lopez et al., 2018). See A.3 for training and implementation details.

As an initial visual assessment of the methods, we compared the UMAPs of PCA and scVI Linear on the same perturbations used in Figures 3 and 4. The UMAPs of both models are shown in Figure A.4 and A.5; both models displayed mixing of the perturbed and unperturbed cells, but better separation than the pre-trained model.

Next, we compared the models on the metrics described in Section 3.4. To ensure fair comparison across methods, we normalised embeddings and distance metrics as described in Section 3.4. The control distributions for various distance measures across different models (Supplementary Figure A.6) highlight the necessity of normalisation for a fair comparison. The contrastive model has the highest median across all 3 distance metrics in the case of separating perturbed from unperturbed transcriptomes (Figure 5, Table 1). PCA proved a competitive baseline on the E-distances, although performing worse than the contrastive model (E-distance medians: PCA= 49.4 vs finetuned= 84.14). In contrast, the pre-trained model without fine-tuning and the highly variable genes showed relatively low performance across all metrics, suggesting limited ability to separate cells based on perturbation-induced transcriptomic changes. The scVI Linear model was the only model to outperform the contrastive model on any of the metrics: while the contrastive model was superior on the Wasserstein distances, scVI Linear had a higher E-distance and Cosine E-distance between perturbation pairs (E-distance medians: scVI Linear = 137.9 vs fine-tuned= 102.0). For all metrics the contrastive model exhibits the largest positive skew, suggesting that it can separate certain perturbations extremely well.

**Table 1:** The medians (over all perturbations in the test set) of different normalized distances in the distributions of perturbed vs unperturbed and perturbation pairs across various models.

| Model | Perturbed vs Unperturbed |  |  | Perturbation Pairs |  |  |
| --- | --- | --- | --- | --- | --- | --- |
|  | E-Dist | Cosine E-Dist | Wass Dist | E-Dist | Cosine E-Dist | Wass Dist |
| Fine-tuned | **84.14** | **74.10** | **29.18** | 102.04 | 103.10 | **37.79** |
| scVI Linear | 76.09 | 68.73 | 11.76 | **137.85** | **124.67** | 15.58 |
| scVI 5L | 42.57 | 38.40 | 19.02 | 76.09 | 66.47 | 25.24 |
| PCA | 49.41 | 45.98 | 9.00 | 83.36 | 75.00 | 9.80 |
| HV | 48.27 | 43.13 | 5.77 | 76.02 | 71.31 | 10.41 |
| Pre-Trained | 10.73 | 8.54 | 9.70 | 17.30 | 14.24 | 14.73 |

**Figure 5.**
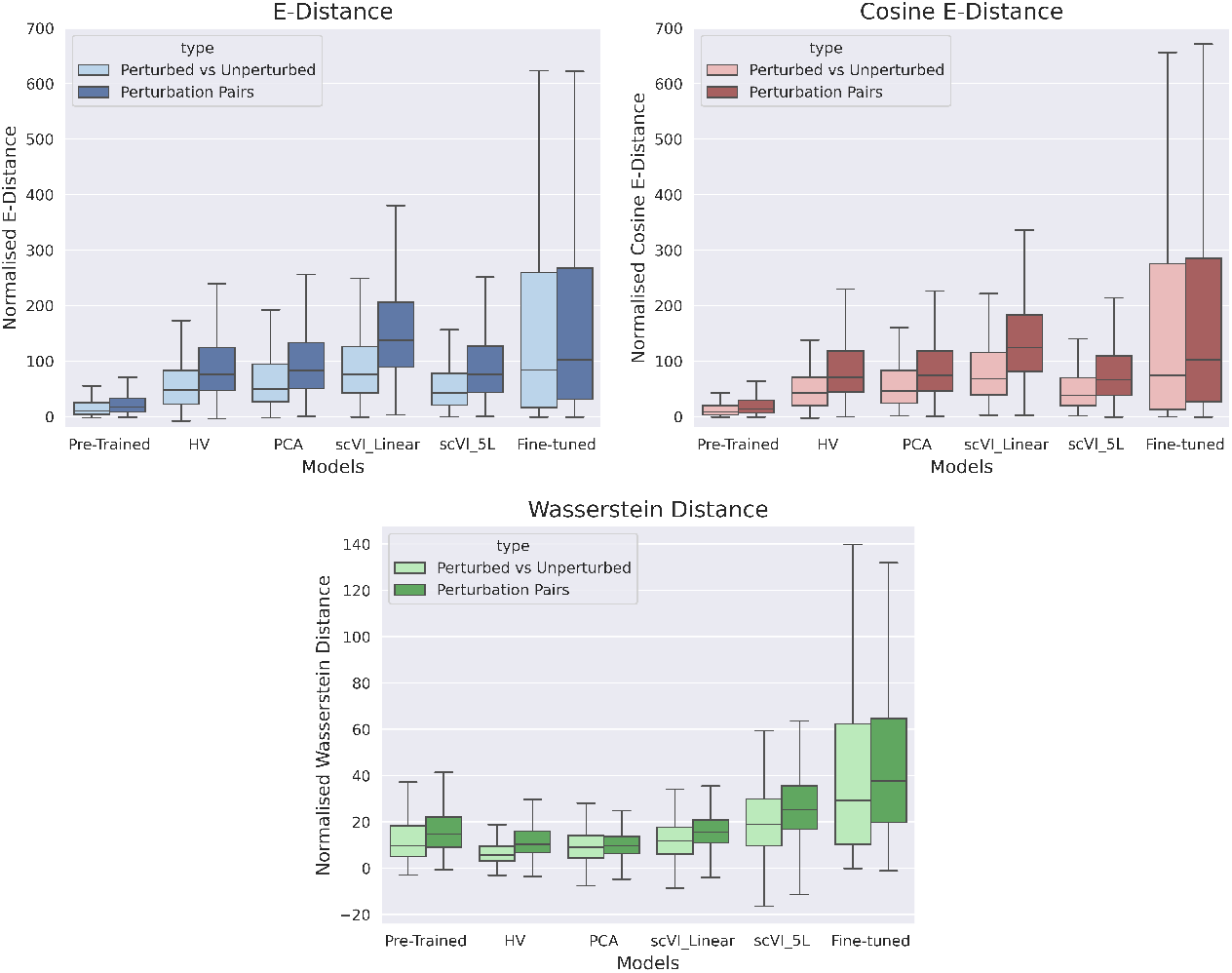
Distance distributions for perturbations in the test set across models. Shown are distances between embeddings of perturbed and unperturbed cells (light colour, *N* = 232 perturbations) and between perturbed cells (dark colour, *N* = 232^2^ perturbation pairs).

These results demonstrate that the contrastive model effectively captures the differences in distributions between perturbed and unperturbed cell profiles, as well as variations within different perturbations. However, scVI Linear exhibited greater separation between perturbation pairs, as indicated by the larger median E-distances, likely due to its ability to leverage latent representations. The high variation of E-distances and Cosine E-distances in the contrastive model suggests that while it effectively distinguishes certain perturbations, others are not optimally separated.

### 4.3 Qualitative assessment of predicted large transcriptomic changes for cells without differentially expressed genes

The model was fine-tuned only on perturbations for which at least 20 DEGs were identified, as we could assume enough signal in this data for the model to learn distinguishing patterns between perturbed and unperturbed cells. Of real interest, however, is whether the model can also be used to identify perturbations that elicited a global transcriptomic change even though no DEGs could be called. To this end, we set out to assess the quality of the contrastive model’s embeddings for perturbed transcriptomes that (i) were excluded from fine-tuning due to having 0 DEGs and (ii) exhibited a high distance metric compared to unperturbed cells in the embedding space of the contrastive model.

In the absence of ground truth, we cannot be certain whether the model detected a global transcriptomic shift for these perturbations or whether the large distances were an artefact of the model. To validate perturbed transcriptomes where the model detected substantial transcriptomic shifts, we examined whether perturbation targets were enriched for gene sets that are particularly prone to inducing significant transcriptional changes when targeted for down-regulation. We focused on genes with critical roles in cellular function. For example, cell cycle genes regulate important processes such as cell growth, DNA replication, and division, ensuring proper cell proliferation and genomic integrity. Similarly, essential genes are indispensable for cell survival and fundamental biological functions, with their disruption typically leading to cell death or failure of vital processes.

To this end, we obtained a list of 663 cell cycle genes (The Gene Ontology Consortium et al., 2023; Ashburner et al., 2000) and 2, 058 essential genes (Replogle et al., 2022). We extracted the cell embeddings of 6, 248 perturbations with 0 DEGs (which were excluded during fine-tuning), obtained embeddings of the same transcriptomes from PCA and scVI Linear for comparison, and ranked the perturbations by their normalised distances from unperturbed cells (from the test set). Regardless of the distance metric used, the contrastive model placed perturbations targeting cell cycle and essential genes more often into the top *n* most distant embeddings than PCA or scVI Linear for different values of *n* (Supplementary Figure A.7). To evaluate the significance of the enrichment, we conducted a permutation test (with 10, 000 permutations) for the top *n* perturbations (according to their E-distances), comparing them to randomly sampled perturbations for different values of *n* (Figure 6). There was a significant enrichment (p-value *<* 0.001, Bonferroni correction for multiple tests) for cell cycle genes for all *n* ≤ 2, 500 and for essential genes for all *n* ≤ 2, 000 . Results were similar when using the two other distance metrics (Supplementary Figure A.8).

**Figure 6.**
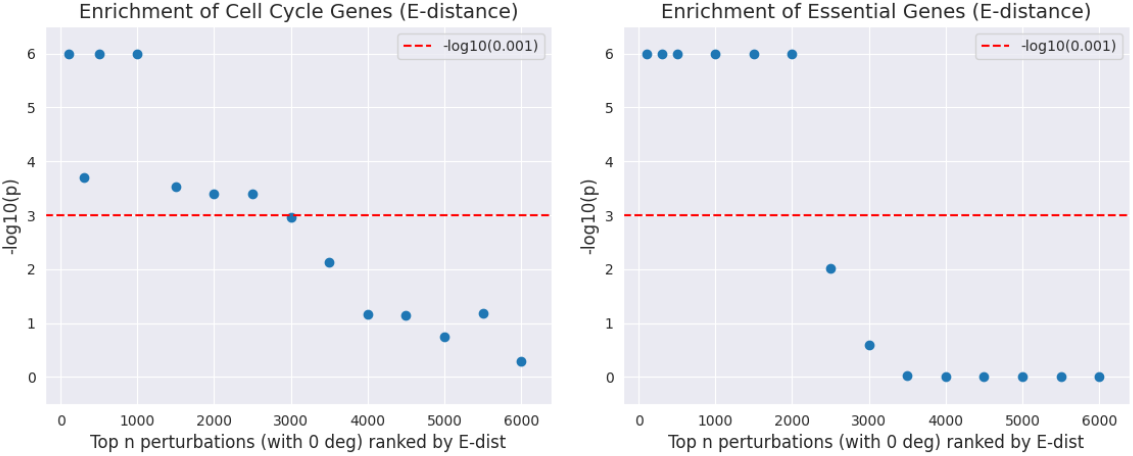
Enrichment analysis of cell cycle (left) and essential genes (right) for perturbations with 0 DEG ranked by normalised E-distance based on the embeddings from the contrastive model. Shown is the log-transformed enrichment p-value for different numbers of top *n* perturbations.

In summary, our contrastive model was able to identify biologically relevant perturbations, even when the number of DEGs failed to capture their effects.

## 5 Discussion

We introduced a novel fine-tuning strategy for single-cell foundation models on Perturb-seq data that leverages contrastive learning to capture transcriptome-level changes in the cell embeddings. We demonstrated the performance of our approach for perturbation analysis by benchmarking our finetuned model against existing approaches using three distance metrics. We focused our comparisons on distance metrics instead of additional clustering as it allows a direct comparison of the separation of the embeddings and ranking of perturbations, independent of the clustering techniques used. We showed that our model identified perturbations that would have been overlooked by traditional differential expression analysis, showing significant enrichment in biologically relevant pathways and functions, including cell cycle regulation and gene essentiality.

In future work, we will explore other contrastive loss functions (Oord et al., 2019) and hybrid methods that integrate contrastive and cross-entropy losses on perturbation targets (Gunel et al., 2021). While our current model was trained on high-DEG perturbations, we will experiment with expanding training to cells with low-DEG perturbations, and evaluate the impact on predictive performance. To mitigate signal dilution, we will explore self-supervised learning approaches that do not rely on predefined similarity labels to learn transcriptome representations (Kim et al., 2021). Further, we will explore explainability techniques developed for BERT-based models to identify genes driving the most significant transcriptomic shifts (Aken et al., 2020; Talebi et al., 2024).

In summary, our method allows for a more comprehensive analysis of perturbation data, thus aiding the identification of perturbations that induce significant transcriptomic changes, and, ultimately, the understanding of disease mechanism.

A Appendix

### A.1 Implementation

Hyperparameters were chosen to allow for distributed learning: max learning rate, 1 × 10^*−*3^ scaled by the number of GPUs; a learning scheduler, linear with warm-up (10k steps) and linear decay; Adam optimizer with weight decay parameter 0.001. The training was distributed over 4 GPUs in one node with a minibatch size 11 and 2 gradient accumulation steps.

To speed up pre-training we used dynamic padding combined with a length-grouped sampler to minimise computation on padding. This sampler takes a randomly sampled megabatch and then orders minibatches by their length in descending order. Mini-batches are then dynamically padded, minimising the computation wasted on padding as sequences of similar lengths are grouped. The authors of Geneformer extended an existing version of this sampler from Huggingface transformers for the distributed case (Theodoris et al., 2023; Wolf et al., 2020). However, neither of these samplers shuffle the mini-batches within the megabatch before passing them to the model, which resulted in a 60x-performance-drop of the trained model in our tests (in terms of training and test perplexity on smaller sample datasets) compared to model runs not employing the grouped-length batching. We implemented a shuffling of the mini batches which slightly diminishes the speed up during training.

For efficient data parallelisation across the GPUS, we used Deepspeed (Rasley et al., 2020). Overall, pre-training was achieved in just over 7 days distributed across one node with four Nvidia A10G 24GB GPUs.

#### A.2 Pre-training evaluation

To compare our pre-trained model to Geneformer (Theodoris et al., 2023), we evaluated both models on a dataset of ∼ 66k peripheral blood mononuclear cells (PBMCs) that was published after both models were trained (Heimlich et al., 2024).

We computed macro-averaged hits@k metrics on masked tokens at different thresholds in the 2000 highest expressed genes in 10k randomly sampled PBMCs (Figure A.1 A). Macro averaging gives equal weight to each gene when computing the accuracy, giving a sense of the model’s performance overall. Here, an instance is one prediction instance, e.g. one of the masked genes in the input we ask the model to fill in. If one gene occurs much more often among the first 2000 genes of the input sequence and is, therefore, more often masked, a model could “cheat” overall metrics by always predicting that one gene. Macro-averaging per gene combats this bias. Both models performed similarly on this task.

To assess further the performance of the models, we followed Kedzierska et al. (2023) and investigated how well the models can reproduce the correct gene rankings per cell given the masked input. To do this, we compare the ground truth order with the predicted order of genes and compute the Spearman correlation coefficients for different cut-offs in all ∼ 66k PBMCs. Similar to Kedzierska et al. (2023), we find that both models struggle to correctly predict the positioning for lower-expressed genes, with our model performing better on reproducing the input rankings of the higher-expressed genes (Figure A.1 B).

#### A.3 implementation and training of scvi

We used the implementation of scVI from *scvi-tools* (Gayoso et al., 2022) and trained a linear VAE with a single encoder and decoder layer (denoted in the text by “scVI Linear”) and a model where both encoder and decoder consisted of 5 hidden layers (denoted in the text by “scVI 5L”). The training data contained all unperturbed and perturbed cells from the training dataset of the contrastive model. The raw scRNA-seq counts of the cells without any further filtering or preprocessing formed the input to the models. We used a random 90/10 split for training and test sets to monitor model convergence. We used a Zero-Inflated Negative Binomial as the likelihood function, and hyperparameters for both models were set at *n latent* = 300, *dropout rate* = 0.1, and *max epochs* = 500 with early stopping.

#### A.4 Supplementary figures

**Figure A1:**
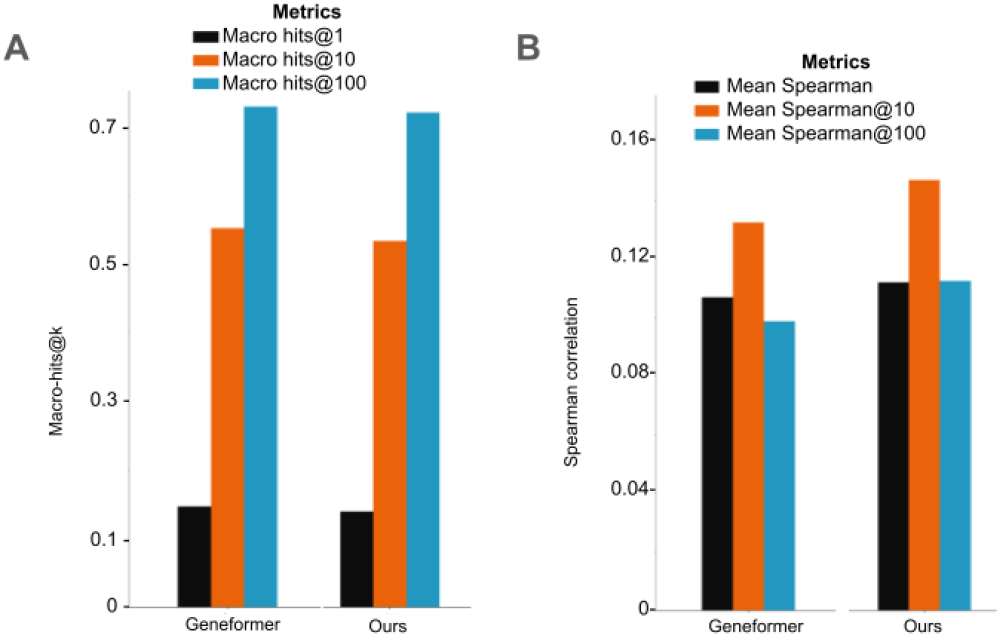
A: Macro-averaged hits@k metrics for *k* = 1, 10 and 100 measuring the performance of correct masked token prediction of the models in 15% masked genes in 10k randomly sampled PBMCs. B: Mean Spearman correlations at different thresholds between ground-truth and predicted gene rankings for the top 100, 500 and 2000 expressed genes per cell in 66k PBMCs. 15% of the input genes were masked at random and the model was asked to generate full ranking outputs for all positions.

**Figure A2:**
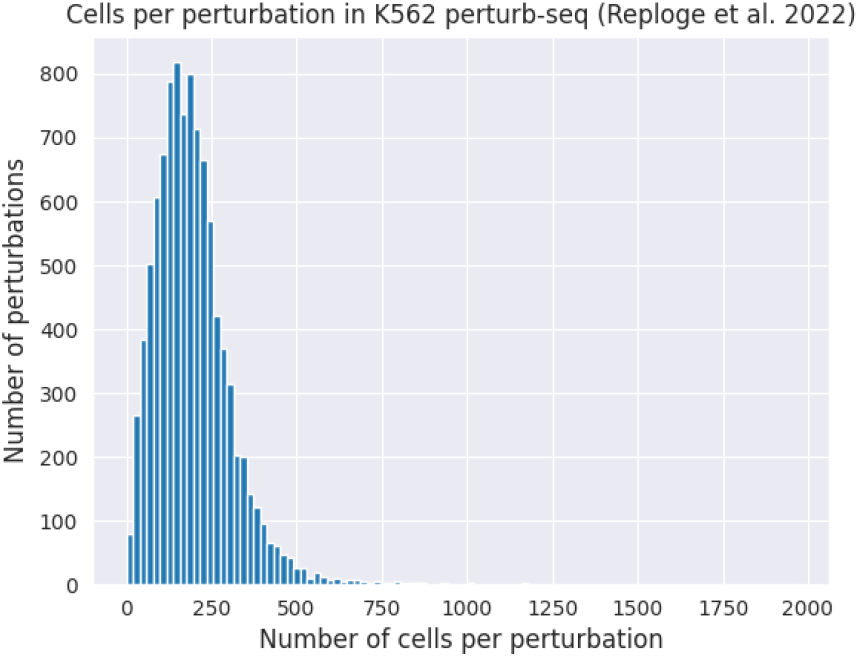
Histogram of the number of cells per perturbation.

**Figure A3:**
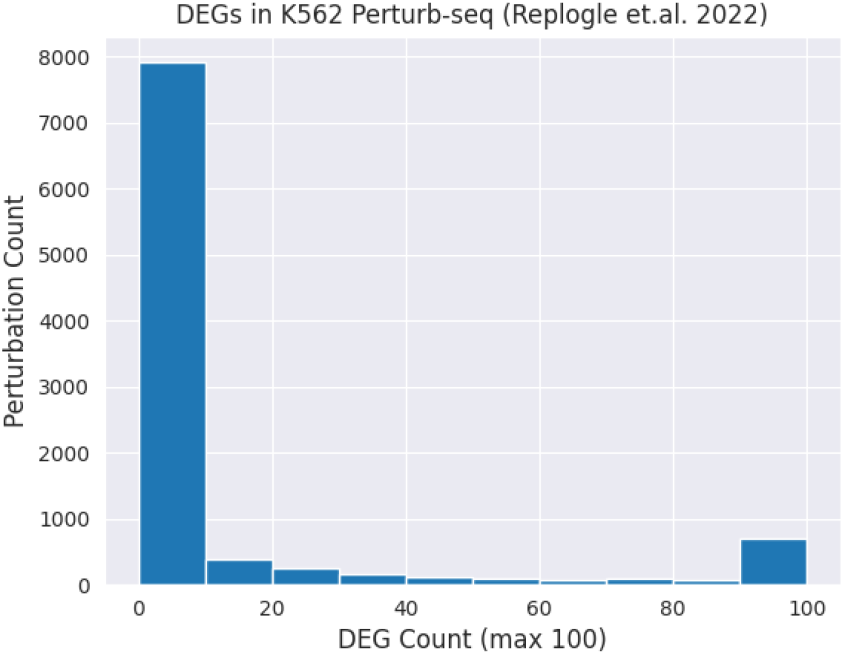
Histogram of the number of DEGs per perturbation. Perturbations with more than 100 DEGs are captured in the last bin.

**Figure A4:**
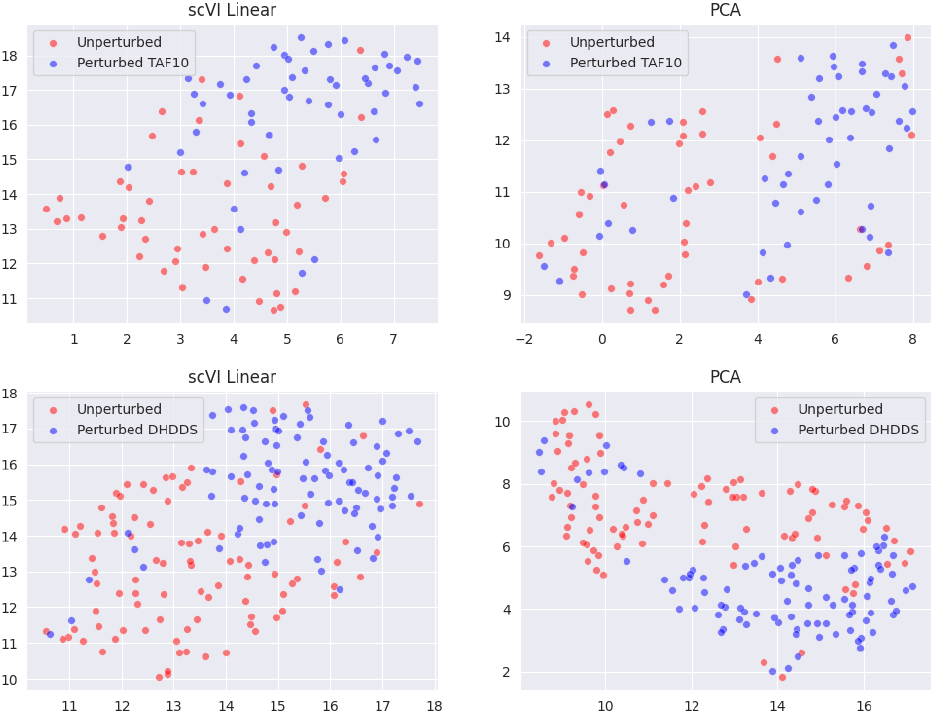
UMAP visualisation of embeddings of perturbed (blue) and unperturbed (red) cells from the test set for the perturbation targets *TAF10* (top) and *DHDDS* (bottom), from PCA (left) and scVI Linear (right).

**Figure A5:**
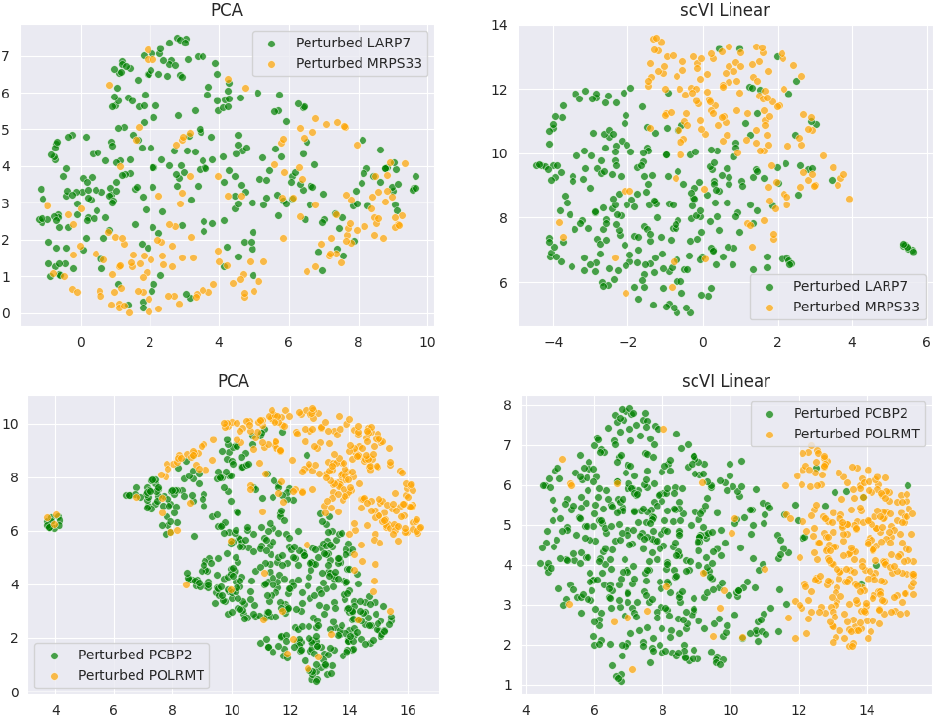
UMAP visualisation of embeddings of perturbed cells from the test set with perturbation targets *LARP7* vs *MRPS33* (top) and *PCBP2* vs *POLRMT* (bottom), from PCA (left) and scVI Linear (right).

**Figure A6:**
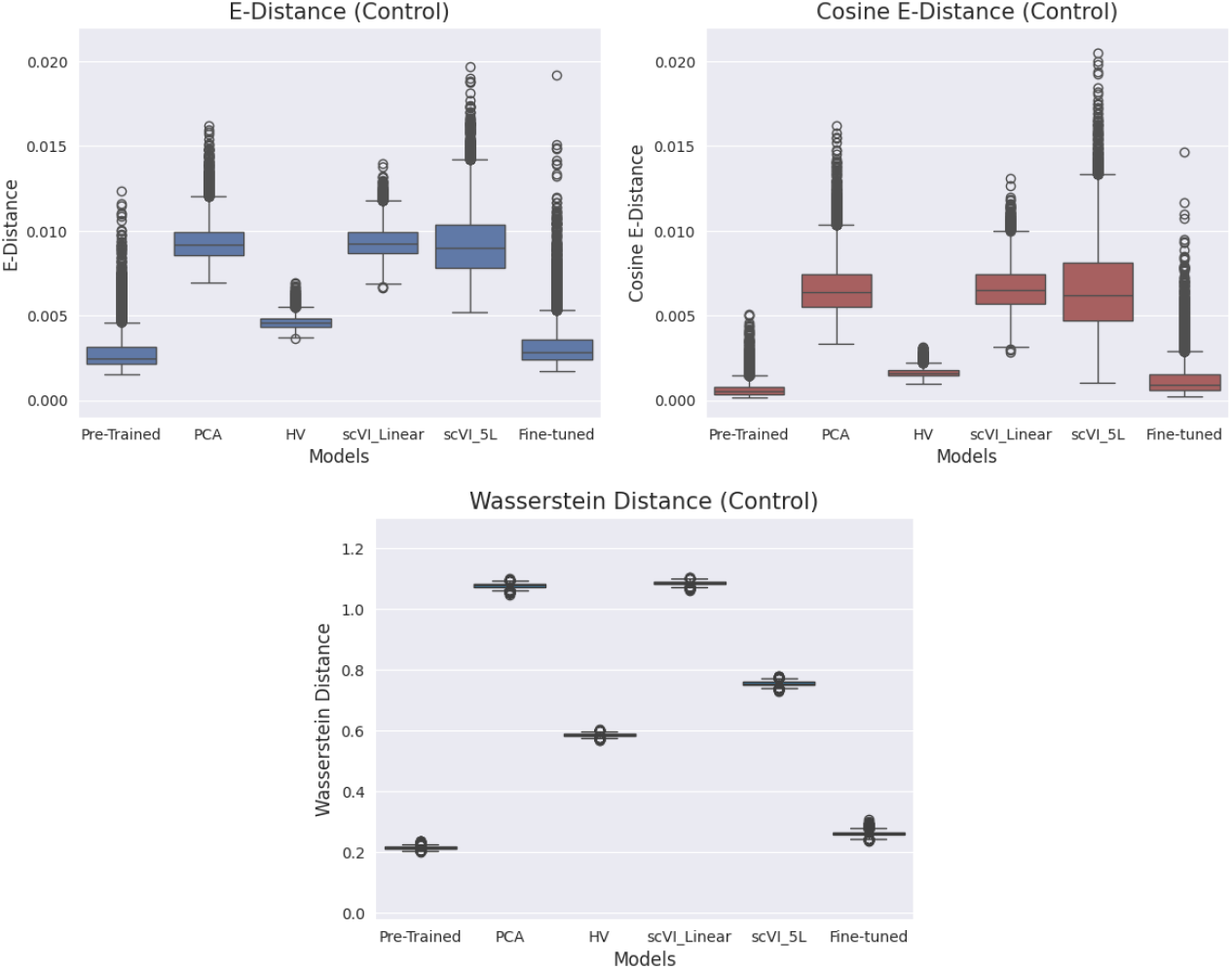
E-distance (top left), Cosine E-distance (top right) and Wasserstein distance (bottom) control distributions of various models. Each data point represents the distance between the embeddings of two groups of randomly sampled unperturbed cells. In total, 10,000 random pairs of groups, each containing 300 cells, were sampled. The difference in control distributions of the models highlights the need to perform z-normalisation to provide fair comparisons across models.

**Figure A7:**
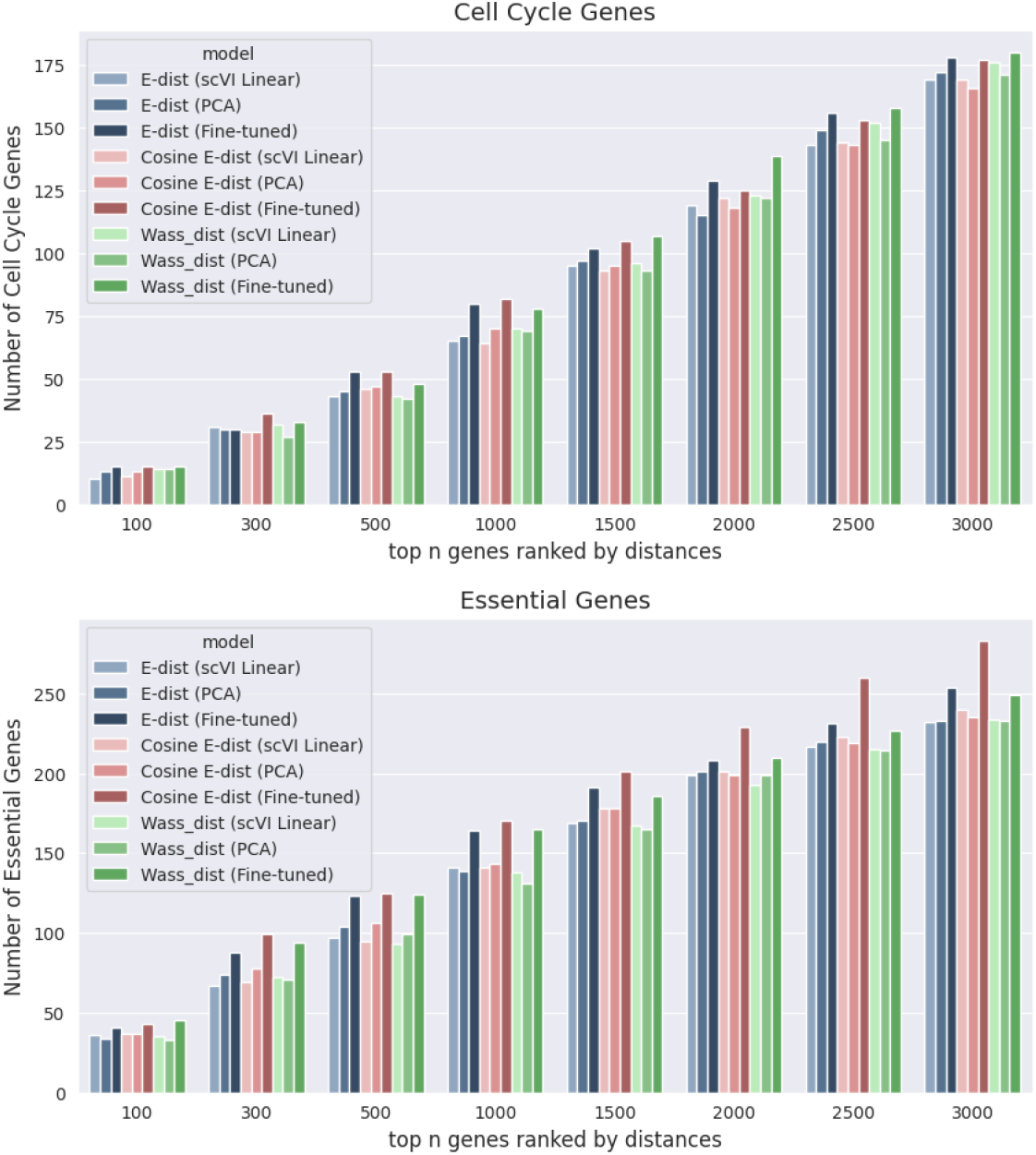
Barchart showing the number of top *n* perturbations ranked by normalised E-distance (blue), normalised cosine E-distance (red) and Wasserstein distance (green) that are targeting either cell cycle genes (top) or essential genes (bottom). Comparing results from scVI Linear (lighter colours), PCA (median colours) and the contrastively fine-tuned model (darker colours).

**Figure A8:**
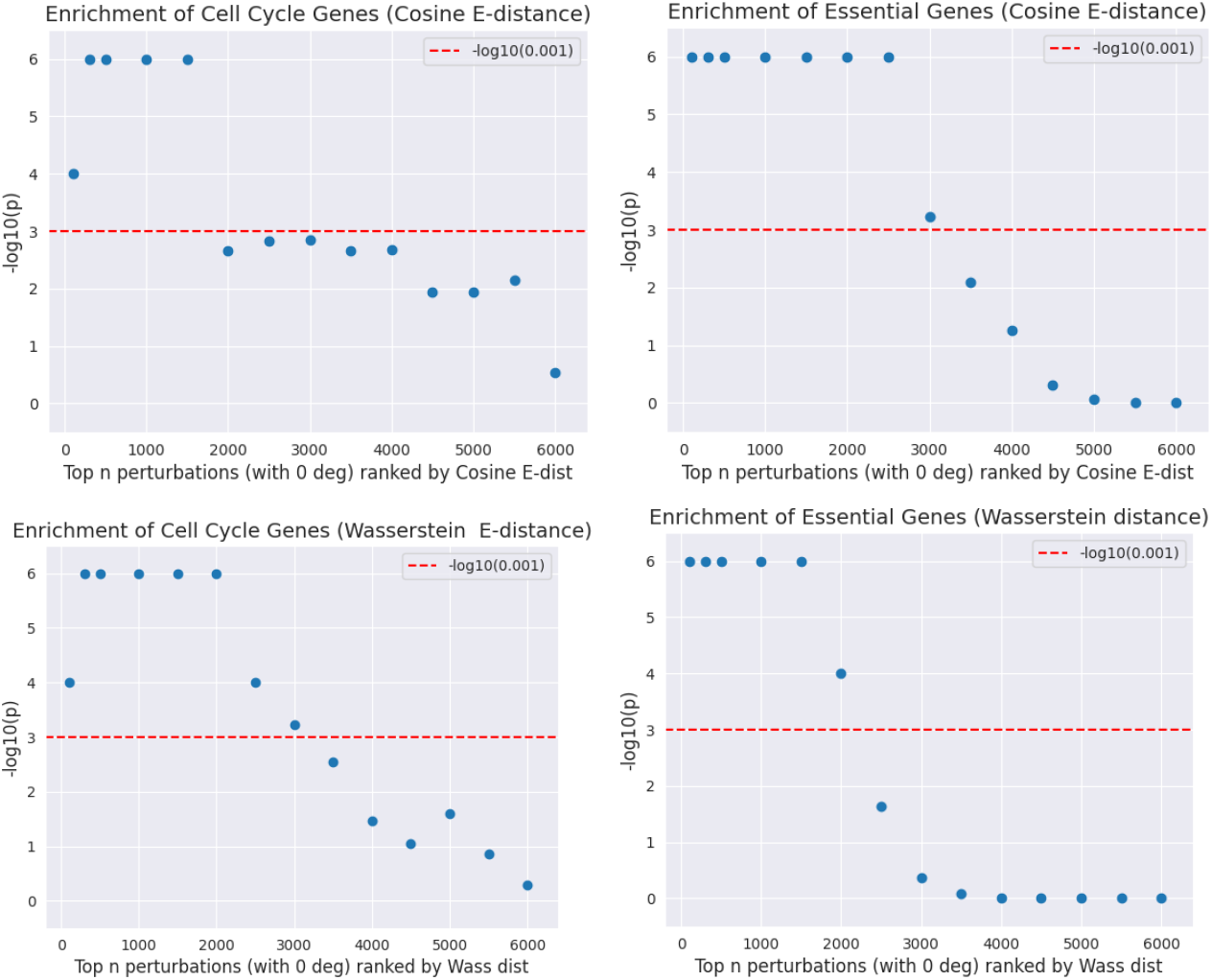
Enrichment analysis of cell cycle (left) and essential genes (right) amongst perturbations with 0 DEGs ranked by normalised Cosine E-distance (top) and normalised Wasserstein distance (bottom) based on embeddings of the contrastive model. The minimum p-value is capped at 1*e* − 6 for the purpose of plotting.

